# Aging alters envelope representations of speech-like sounds in the inferior colliculus

**DOI:** 10.1101/266650

**Authors:** Aravindakshan Parthasarathy, Björn Herrmann, Edward L. Bartlett

## Abstract

Hearing impairment in aging is thought to arise from impaired temporal processing in auditory circuits. We used a systems-level (scalp recordings) and a microcircuit-level (extracellular recordings) approach to investigate how aging affects the sensitivity to temporal envelopes of speech-like sounds in rats. Scalp-recorded potentials suggest an age-related increase in sensitivity to temporal regularity along the ascending auditory pathway. The underlying cellular changes in the midbrain were examined using extracellular recordings from inferior colliculus neurons. We observed an age-related increase in sensitivity to the sound’s onset and temporal regularity (i.e., periodicity envelope) in the spiking output of inferior colliculus neurons, relative to their synaptic inputs (local field potentials). This relative enhancement for aged animals was most prominent for multi-unit (relative to single-unit) spiking activity. Spontaneous multi-unit, but not single-unit, activity was also enhanced in aged compared to young animals. Our results suggest that aging is associated with altered sensitivity to a sound’s temporal regularities, and that these effects may be due to increased gain of neural network activity in the midbrain.

## INTRODUCTION

Older and middle-aged listeners experience difficulties understanding speech, particularly in challenging listening situations such as in the presence of background sounds, competing speakers, or reverberation (Ruggles et al., 2011, Pichora-Fuller and Souza, 2001). Speech perception depends on the sensitivity of the auditory system to temporal regularities in sounds. Temporal regularities in speech may be divided into three categories: the slow fluctuations (<50 Hz) of the speech envelope that capture word and syllabic rate; the periodicity envelope (50–500 Hz), which contains the fundamental frequency of the speaker’s voice (f0) and is crucial for speaker identification (Bregman, 1990); and the temporal fine structure (>500 Hz), which contains information about formant structure (Rosen, 1992). The sensitivity of the auditory system to temporal regularity in sounds declines with age, with drastic consequences for speech perception (Walton, 2010, Fullgrabe et al., 2015, Anderson et al., 2011), but the neurophysiological changes that underlie this age-related decline are not well understood.

An age-related decline in temporal processing abilities is thought to be primarily neural in origin, because it is independent of changes in hearing thresholds due to impaired cochlear function (Frisina and Frisina, 1997, Pichora-Fuller and Souza, 2001, Gordon-Salant and Fitzgibbons, 2001). The neural deficits that may contribute to impaired sensitivity to temporal regularity include cochlear synaptopathy – that is, the degradation of cochlear synapses between inner hair cells and auditory nerve fibers (Sergeyenko et al., 2013) – and a decrease in inhibitory neurotransmitters in the brainstem, midbrain and cortex (Caspary et al., 2008, Takesian et. al., 2009, Rabang et al., 2012). Loss of inhibition may result in increased neural activity in central auditory regions despite diminished inputs from peripheral structures (Mohrle et al., 2016, Parthasarathy et al., 2014, Hughes et al., 2010). However, it is less clear how aging affects sensitivity to temporal regularity in central auditory regions, in particular for complex, speech-like sounds.

Previous work suggests that neurons in the inferior colliculus show altered temporal processing including changes in the sensitivity to the temporal regularities in sounds (Walton et al., 2002, Walton et al., 1998, Palombi et al., 2001, Rabang et al., 2012, Schatteman et al., 2008). These studies focused on neuronal spiking, which reflects the output of neurons. In contrast, local field potentials (LFPs) are thought to reflect the summed synaptic inputs to a neuron or local neuronal population to a large degree (but other non-synaptic activity might additionally contribute; (Bullock, 1997, Buzsaki et al., 2012, Logothetis and Wandell, 2004, Logothetis et al., 2001)). Recent recordings of LFPs and spiking activity show that synchronization of spiking activity (output) is abnormally enhanced in the aging inferior colliculus, despite decreased synaptic inputs (input), and that this age-related relative increase in activity (i.e., from a neuron’s input to its output) is specific for sounds with modulation rates below 100 Hz (Herrmann et al., 2017). Whether aging also leads to an over-sensitivity to temporal regularities in complex, speech-like sounds is unknown.

In the current study, we test the hypothesis that neural synchronization to the periodicity envelope (~100 Hz) of speech is abnormally enhanced in the inferior colliculus of aged animals. We assess peripheral neural function by measuring wave 1 amplitudes of the auditory brainstem responses (ABRs) and show physiological evidence for cochlear synaptopathy in aged animals. Scalp-recorded neural synchronization to the envelope of a speech-like stimulus is increased specifically in more rostral regions in the auditory pathway such as the inferior colliculus compared to more caudal ones such as the auditory nerve. Further, we assess the relation between LFPs (synaptic input) and spiking output in the inferior colliculus using extracellular recordings. Synchronization of LFPs to the envelope of a speech-like sound is decreased in aged animals, whereas synchronization of spiking activity from well isolated units does not differ between age groups. Multi-unit spike synchronization, however, is drastically increased for aged animals, suggesting changes in gain control mechanisms occurring largely at a neural network level.

## METHODS AND MATERIALS

### Ethical approval

The experimental procedures described in the present investigation were approved by the Institutional Animal Care and Use Committee of Purdue University (PACUC #1111000167). The experiments included in this study comply with the policies and regulations described by (Drummond, 2009). Rats were housed one per cage in accredited facilities (Association for the Assessment and Accreditation of Laboratory Animal Care) with food and water provided *ad libitum*. The number of animals used was reduced to the minimum necessary to allow adequate statistical analyses.

### Subject population

The study design is cross-sectional and used male Fischer-344 rats obtained from Charles River Laboratories and Taconic. The aging Fisher-344 rat has been suggested to be a suitable model to study presbycusis in aging animals (Syka, 2010) and has been shown to exhibit high frequency hearing loss and metabolic presbycusis that are characteristic of human age-related loss of hearing sensitivity (Dubno et al., 2013, Allen and Eddins, 2010). 11 young (3–6 months, ~300 g) and 12 aged (22–26 months, ~400– 500 g) animals were used for scalp recordings, and 11 young (3–6 months, ~300 g) and 9 aged (22–26 months, ~400–500 g) animals were used for extracellular recordings. A subset of the ABR recordings have been reported in previous studies (Parthasarathy and Bartlett, 2012, Parthasarathy et. al., 2014).

### Sound stimulation

The stimulus was a natural English syllable, /ba/, which was 260ms long and spoken by a male speaker of North American English with a fundamental frequency ~110Hz. In order to account for the differences in the hearing range between rats and humans, as well as to increase the number of responsive neurons in the inferior colliculus of rats, this speech token was half-wave rectified, and used to modulate a broadband noise carrier (0.04–40 kHz) (Figure 1A). This preserved the periodicity envelope as well as the original fine stricture of the speech token, both of which served as the modulator for the broadband noise (Figure 1B, C). This stimulus design was chosen to understand how neurons represent temporal regularities that are important for human speech and to aid comparison with human literature, rather than investigating the representation of conspecific vocalizations that may not generalize to human speech processing.

**Figure 1:**
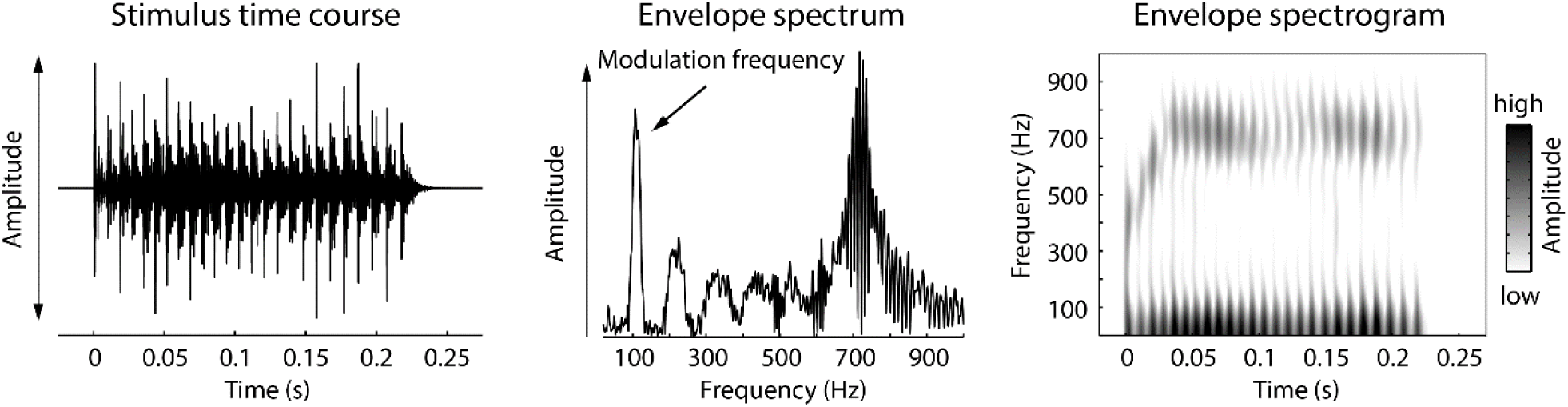
Speech-like sound used in the current study. **Left:** Waveform of the speech-like sound. **Middle:** Amplitude spectrum of the envelope of the speech-like sound displayed on the left. **Right:** Spectrogram of the envelope of the speech-like sound.

### Stimulus generation

Sound stimuli were generated using SigGenRP (Tucker-Davis Technologies, TDT) sampled at a rate of 97.64 kHz (standard TDT sampling rate) and presented through custom-written interfaces in OpenEx software (TDT). Sound waveforms were generated via a multichannel processor (RX6, TDT), amplified (SA1, TDT), and presented free-field through a Bowers and Wilkins DM601 speaker. The output from the speaker was calibrated free field, using SigCal (TDT) and a Bruel Kjaer microphone with a 0.25-in. condenser, pointed at frontal incidence to the speaker, from the same location as the animal’s right ear, and was found to be within ±6 dB for the frequency range tested. All recordings took place in an Industrial Acoustics booth lined with 1 inch (35 mm) Sonex foam with ~90% absorption for frequencies ≥1000 Hz, minimizing potential echoes or reverberations. All analyses described below accounted for the travel time of the sound wave from the speaker to the animals’ ears.

### Electrophysiological recordings at the scalp

Methods for experimental setup, sound stimulation, and scalp-potential recordings are similar to those described before (Parthasarathy et al., 2016, Parthasarathy et al., 2014). The animals were briefly anesthetized using isoflurane (1.5–2%) for the insertion of subdermal needle electrodes (Ambu) and the intramuscular injection of dexmedetomidine (Dexdomitor, 0.2 mg/kg), an α-adrenergic agonist that acts as a sedative and an analgesic. Recordings commenced after a 15 minute recovery from the anesthesia, and were terminated if the animal showed any signs of discomfort like increased breathing or major movement as monitored by a video camera in the recording chamber.

The scalp-evoked responses to the temporal envelope of the speech-like sound (Envelope following response - EFR) were obtained using two simultaneous recording channels. One positive electrode (caudal channel) was placed on the animal’s forehead along the midline at Cz – Fz. This electrode has a strong wave 3 ABR component (cochlear nuclei) and best sensitivity to modulation frequencies ≥100 Hz (Parthasarathy and Bartlett 2012). Another positive electrode (rostral channel) was placed horizontally, along the inter-aural line, above the location of the inferior colliculus. This electrode has a strong wave 1, 4 and 5 ABR components (auditory nerve, the inferior colliculus and its afferents) and best sensitivity to modulation frequencies <100 Hz (Parthasarathy and Bartlett, 2012). The negative electrode was placed under the right ear, along the mastoid, and the ground was placed in the nape of the neck. Impedances were ensured to be always less than 1KΩ as tested using the low-impedance amplifier (RA4LI, TDT).

This two-channel setup allowed greater sensitivity to different ranges of modulation frequencies and putative generators compared to recording from a single channel alone, as reported previously (Parthasarathy and Bartlett, 2012). Signal presentation and acquisition were performed by BioSig software (TDT). The stimulus was presented free field to the right of the animal, at a distance of 115 cm from speaker to the right ear. Digitized waveforms were recorded with a multichannel recording and stimulation system (RZ-5, TDT) and analyzed with BioSig or custom written programs in MATLAB (Mathworks).

### Surgical procedures for extracellular recordings

Methods for surgery, sound stimulation and recording are similar to those described in (Herrmann et al., 2015, Rabang et al., 2012). Surgeries and recordings were performed in a 9’×9’ double walled acoustic chamber (Industrial Acoustics Corporation). Animals were anesthetized using a mixture of ketamine (VetaKet, 80 mg/kg) and dexmedetomidine (Dexdomitor, 0.2 mg/Kg) administered intra-muscularly via injection. A constant physiological body temperature was maintained using a water-circulated heating pad (Gaymar) set at 37°C with the pump placed outside the recording chamber to eliminate audio and electrical interferences. The animals were maintained on oxygen through a manifold. The pulse rate and oxygen saturation were monitored using a pulse-oximeter to ensure they were within normal ranges during surgery. Supplementary doses of anesthesia (20mg/kg of ketamine, 0.05mg/kg of dexmedetomidine) were administered intra-muscularly as required to maintain areflexia and a surgical plane of anesthesia. An initial dose of dexamethasone and atropine was administered prior to incision to reduce swelling and mucosal secretions. A subdermal injection of Lidocaine (0.5 ml) was administered at the site prior to first incision. A central incision was made along the midline, and the calvaria exposed. A stainless steel headpost was secured anterior to bregma using an adhesive and three screws drilled into the skull to provide structural support for a head-cap, constructed of orthodontic resin (Dentsply). A craniotomy was performed from 9–13 mm posterior to bregma, which extended posterior to the lambda suture, and 3 mm wide extending from the midline. The dura mater was kept intact, and the site of recording was estimated stereotaxically using a rat atlas (Paxinos and Watson, 2006) as well as using internal vasculature landmarks and physiological measurements. At the completion of recordings, animals were euthanized with Beuthanasia (200 mg/kg IP). Once areflexive, they were perfused transcardially with 150-200 mL phosphate buffered saline with followed by 400–500 mL 4% paraformaldehyde. The brain was then removed and stored or processed further for Nissl or immunohistochemistry.

### Electrophysiological recordings in extracellular space

Neural activity in inferior colliculus was recorded *in vivo* using a tungsten microelectrode (A-M Systems) encased in a glass capillary that was advanced using a hydraulic micro-drive (Narishige). We recorded 118 units in young rats and 121 units in aged rats. The inferior colliculus was identified based on short-latency driven responses to tone stimuli. The central nucleus of the inferior colliculus was identified using the ascending tonotopy moving in a dorsoventral direction as well as narrowly tuned responses to pure tones with frequencies ranging from 0.5 to 40 kHz. Recordings were obtained from both the dorsal cortex and central nucleus. Although we cannot exclude that we recorded from dorsal and lateral (or external) cortex, based on the recording depth, the presence of sustained responses to tones and amplitude-modulated stimuli, as well as clear frequency tuning in most cases, we estimate that most of our units were recorded from the central nucleus.

The sounds were presented to the animal at azimuth 0° and elevation 0°. Neural signals were acquired using the tungsten microelectrode connected to a headstage (RA4, TDT) and were amplified (RA4PA preamplifier, TDT). The digitized waveforms and spike times were recorded with a multichannel recording and stimulation system (RZ-5, TDT) at a sampling rate of 24.41 kHz (standard TDT sampling rate). The interface for acquisition and spike sorting were custom made using the OpenEx and RPvdsEx software (TDT). The units acquired were filtered between 500 Hz (occasionally 300 Hz) and 5000 Hz. The acquired spikes were stored in a data tank and analyzed using custom written software in Matlab. Local field potentials were simultaneously recorded from the same electrode by sampling at 1525.88 Hz and bandpass filtering from 3 to 500 Hz.

### Assessment of peripheral and brainstem function

Auditory brainstem responses (ABRs) were recorded using the scalp-recording setup described above. ABRs were recorded in response to brief broadband click stimuli of 0.1-ms duration that varied in sound levels from 5 to 95 dB SPL in 10 dB steps. The stimuli were presented in alternating polarity at 26.6 clicks per second. The acquisition window was 20 ms, and each ABR was an average of 1,500 repetitions. The ABR amplitudes of different waves were calculated as the amplitude of the peak of the wave from the baseline, in BioSig (TDT). ABRs between groups were compared at peak response level, which was determined as the lowest sound level that produced the maximum amplitude for each animal. This typically corresponded to 70–75 dB sound pressure level (SPL) for the young and 80–85 dB SPL for the aged rats.

### Analysis of neural synchronization recorded at the scalp

For scalp recorded neural synchronization (i.e., EFRs), the speech-like stimulus (described above) was presented with a repetition rate of 3.1 Hz. The stimulus was presented at peak response level described above, which was determined following a fast-Fourier transform (FFT) of the time-domain response. Each response time course was obtained as an average of 200 stimulus repetitions in alternating polarity. Responses were filtered online between 30–3000 Hz with a 60 Hz notch filter.

In order to analyze the fidelity of the auditory system to synchronize with the temporal structure in the speech-like sound, we first used a broad-scale approach by calculating the correlation between the stimulus waveform and the response time course for lags ranging from 0 to 0.05 s. This cross-correlation approach was calculated twice, once for a low-frequency range (i.e., stimulus waveform and response time course were low-pass filtered at 300 Hz; Butterworth) and once for a high-frequency range (i.e., stimulus waveform and response time course were band-pass filtered from 300 to 3000 Hz; Butterworth). The former assessed neural synchronization to the vowel-like envelope periodicity, the latter assessed neural synchronization to the temporal fine structure in the stimulus. The highest correlation value (out of all lags) was used as a measure of synchronization strength. Separately for the two channels and the two filtered signals, Wilcoxon’s rank sum test (Matlab: ranksum) was used to test whether correlation values differed between age groups.

We further investigated neural synchronization by calculating the amplitude spectrum (using a fast Fourier transform [FFT]; Hann window; zero-padding) of the averaged time-domain signal. Neural synchronization could not be quantified as inter-trial phase coherence (Lachaux et al., 1999) because the recordings were averaged across trials online. This analysis focused on neural synchronization to the envelope of the speech-like sound (~110 Hz) and we thus averaged the spectral amplitudes in the frequency window ranging from 105–115 Hz. Wilcoxon’s rank sum test (Matlab: ranksum) was used to test whether neural synchronization to the envelope differed between age groups (separately for the caudal and rostral channel).

### Analysis of local field potentials (LFPs)

Local field potentials previously band-pass filtered between 3-500Hz during acquisition were further notch filtered at 60 Hz and 120 Hz (elliptic filter; infinite-impulse response [IIR}; zero-phase lag) to suppress line noise, and low-pass filtered at 200 Hz (Butterworth; IIR; zero-phase lag).

For the time-domain analysis, single-trial time courses were averaged separately for each age group. Onset responses were analyzed by calculating the root-mean-square (RMS) amplitude of the averaged signal in the 0–0.08 s time window. Wilcoxon’s rank sum test (Matlab: ranksum) was used to test differences in sound-onset responses between age groups.

Neural synchronization was analyzed by calculating a normalized vector strength spectrum (Wolff et al., 2017, Herrmann et al., 2017). To this end, for each trial, a fast Fourier transform (FFT; Hann window; zero-padding) was calculated using the data ranging from 0.08 s to 0.225 s post sound onset (frequency range: 20–180 Hz; step size: 0.05 Hz; zero-padding) to exclude onset and offset transients. An inter-trial phase coherence (ITPC) spectrum was calculated using the complex values from the FFT (Lachaux et al., 1999). An ITPC permutation distribution was generated by flipping the sign of a random subset of trials (Wolff et al., 2017), followed by ITPC calculation. This procedure was repeated 200 times and resulted in a permutation distribution of ITPC values. The spectrum of normalized vector strength was calculated by subtracting the mean ITPC of the permutation distribution from the empirically observed ITPC and dividing the result by the standard deviation of the permutation distribution (separately for each frequency). The normalized ITPC (vector strength) reflects a statistical measure – that is, a z-score – with a meaningful zero that indicates non-synchronized activity.

In order to test for differences in neural synchronization at the envelope frequency between age groups, the normalized vector strength was averaged across the 105–115 Hz frequencies. Wilcoxon’s rank sum test (Matlab: ranksum) was used to test whether neural synchronization differed between age groups.

### Analysis of multi-unit activity (MUA)

Multi-unit activity was extracted based on the recorded broad-band neural signal (Lakatos et al., 2013, Lakatos et al., 2005). To this end, the signal was high-pass filtered at 300 Hz (Butterworth; IIR; zero-phase lag), full-wave rectified by calculating absolute values, low-pass filtered at 200 Hz (Butterworth; IIR; zero-phase lag), and down-sampled to the sampling frequency of the LFP (1525.88 Hz). Data analysis followed closely the analysis of LFP data.

For the time-domain analysis, single-trial time courses were averaged separately for each age group. Onset responses were analyzed by calculating the mean amplitude of the averaged signal in the 0–0.08 s time window. Wilcoxon’s rank sum test (Matlab: ranksum) was used to test differences in sound-onset responses between age groups.

Neural synchronization was analyzed by calculating a normalized vector strength spectrum using the same procedure as described for the LFP data.

### Analysis of single-unit activity (SUA)

Overall, we recorded from 89 single units in young, and 123 single units in the aged animals. Single unit activity was isolated online during the recordings of the neural signals. Units that were substantially above noise threshold were sorted visually online, and then subsequently identified and isolated based on waveform similarity offline, using the OpenEx interface. Isolated single units had an SNR of at least 6 dB relative to the surrounding noise floor and stable waveform shapes. The acquired spikes were stored in data tank and analyzed using custom-written software in MATLAB.

Peri-stimulus time histograms were calculated by convolving an impulse vector generated from spike times with a Gaussian function with a standard deviation of 3 ms (Dayan and Abbott, 2001). Neural responses to the onset of the sound was investigated by calculating the mean firing rate for the 0–0.08 s time window. Wilcoxon’s rank sum test (Matlab: ranksum) was used to test differences in sound-onset responses between age groups.

Investigation of neural synchronization was assessed using normalized vector strength based on spike times (Herrmann et al., 2017). To this end, spike times (within the 0.08–0.225 s time window) were transformed to phase angles (***p***) using the following formula:

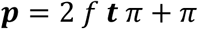

where *f* is frequency and ***t*** a vector of spike times. Phase angles were wrapped to range from –π to π. The empirical vector strength *v* (similar to ITPC described above), that is, the resultant vector length, was calculated as follows (Lachaux *et al.*, 1999):

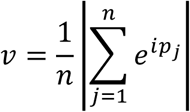

where *v* is the vector strength, *i* the imaginary unit, ***p*** the vector of phase angles, and *n* the number of spikes (with *j* being the index). Vector strength can be biased by the number of spikes, with smaller *v* values for a higher number of spikes. In order to avoid biases estimates, a distribution of random vector strength values was calculated. That is, *n* (number of spikes) random phase values between –π and π were generated and the vector strength (i.e., the resultant vector length) was calculated. Randomly drawing phase values and calculation of the vector strength was repeated 1000 times, which resulted in a distribution of random vector strengths given the number of spikes. The normalized vector strength was then calculated by subtracting the mean of the random vector strength distribution from the empirical vector strength and dividing the result by the standard deviation of the random vector strength distribution. The normalized vector strength was calculated for frequencies (*f*) ranging from 20 Hz to 180 Hz with a frequency resolution of 0.1 Hz, resulting in a vector strength spectrum. In order to test for differences in neural synchronization at the envelope frequency between age groups, the normalized vector strength was averaged across the 105–115 Hz frequencies. Wilcoxon’s rank sum test was used to test whether neural synchronization differed between age groups.

### Effect size measure

Throughout the manuscript, effect sizes are provided as r_e_ (r_equivalent;_ Rosenthal and Rubin, 2003). r_e_ is equivalent to a Pearson product-moment correlation for two continuous variables, to a point-biserial correlation for one continuous and one dichotomous variable, and to the square root of partial η^2^ for an analysis of variance.

## RESULTS

### ABR amplitudes are reduced in aged animals

ABRs were recorded simultaneously from two electrode montages – one that emphasizes more caudal generators (putatively including the auditory nerve and the cochlear nucleus), and another that emphasizes more rostral generators (putatively including the inferior colliculus), as evidenced by the differences in ABR waveform morphology, modulation rate sensitivity, and the effects of anesthesia (Parthasarathy and Bartlett, 2012, Parthasarathy et al., 2014).

ABR wave 1 amplitudes were significantly decreased for aged compared to young rats (Figure 2A, rostral channel: p = 1.96^e-4^, r_e_ = 0.701, df = 21; caudal channel: p = 7.20^e-5^, r_e_ = 0.732, df = 21). The wave 1 of the ABR originates in the auditory nerve, and its amplitude at suprathreshold sound levels is a physiological indicator for the degree of cochlear synaptopathy due to aging or noise exposure (Sergeyenko et al., 2013, Stamper and Johnson, 2015). This age-related decrease in ABR amplitudes was also observed for all subsequent waves (wave 3: p = 5.55^e-5^, r_e_ = 0.739, df = 21; wave 4: p = 9.92^e-4^, r_e_ = 0.641, df = 21; wave 5: p = 7.17^e-5^, r_e_ = 0.732, df = 21; Figure 2A) with generators in other brainstem and midbrain nuclei. Hence, there was an overall reduction in transient responses in the auditory nerve fibers, auditory brainstem and midbrain with age.

**Figure 2:**
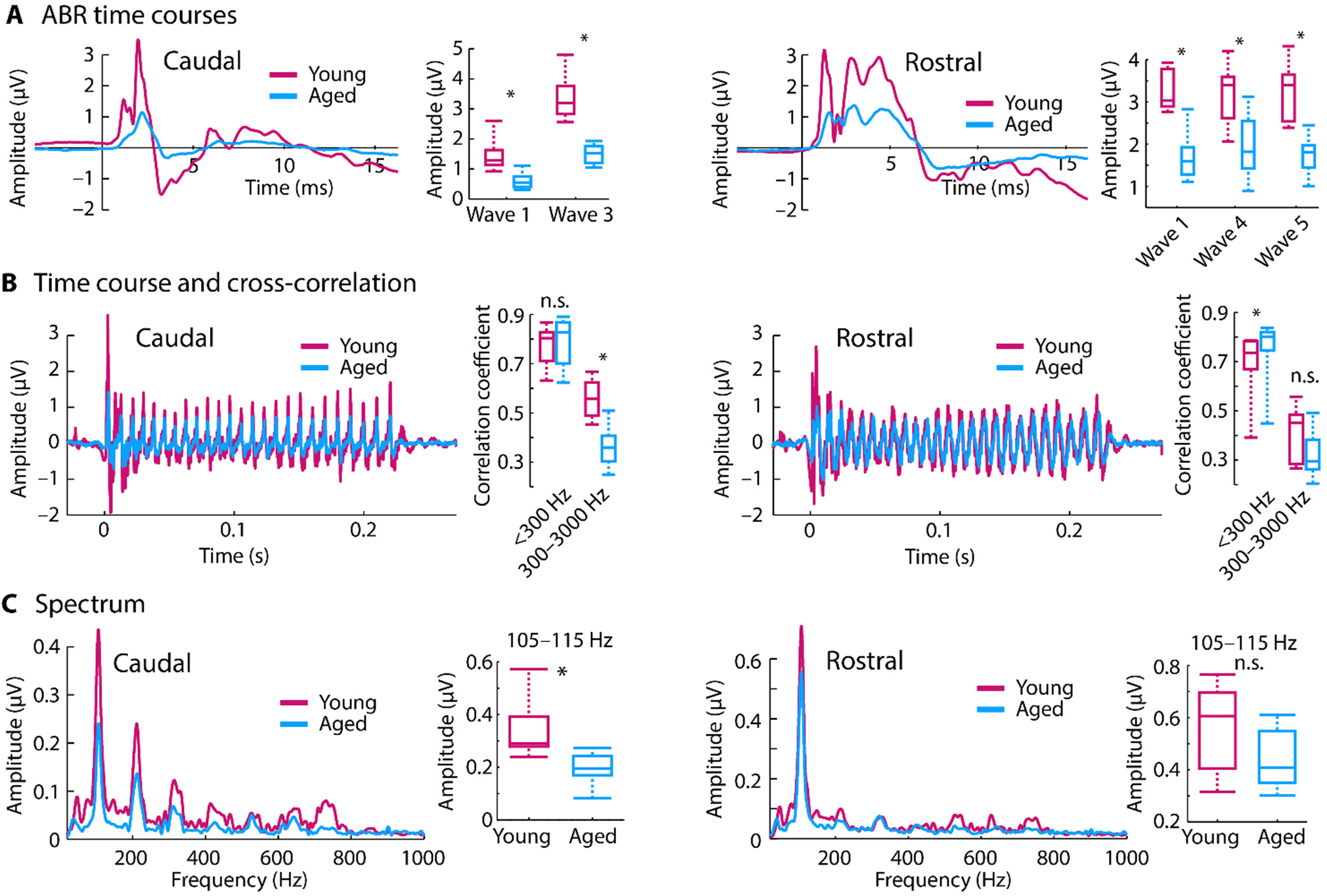
Scalp-recorded potentials simultaneously recorded in two channels emphasizing rostral versus caudal generators. **A:** Averaged click-evoked auditory brainstem responses (ABRs) and amplitudes for different ABR waves. **B:** Time courses of envelope following responses (EFRs) in response to the speech-like /ba/ sound. Boxplots show coefficients from the cross-correlation for different frequency bands. **C:** Amplitude spectra derived from fast Fourier transforms. Boxplots show neural synchronization strength to the sound’s F0 envelope (105–115 Hz). *p < 0.05, n.s. – not significant

### Scalp-recorded neural synchronization shows strong spectral peaks and stimulus correlation with age despite weak ABR amplitudes

The ability of the auditory system to represent the various temporal regularities of speech were assessed using a measure of neural synchronization (i.e., EFR) evoked to the speech-like sound. The sensitivity of neural activity to the sound’s temporal structure was assessed in two ways: Cross-correlation and spectral amplitude (derived from a fast Fourier transform). The overall temporal sensitivity was calculated by cross-correlating the time course of the scalp-recorded neural response with the stimulus time course. In the caudal channel, cross-correlation values were reduced for aged compared to young rats (p = 1.54^e-4^, r_e_ = 0.709, df = 21) for the 300–3000-Hz frequency range that contains information about the sound’s temporal fine structure. There was no difference in correlation values between young and aged animals for the <300-Hz frequency range (envelope) (p = 0.479, r_e_ = 0.155, df = 21; Figure 2B, left). In the rostral channel with putative generators in the midbrain and its afferents, there was no age difference for the 300–3000-Hz frequency range (p = 0.069, r_e_ = 0.385, df = 21). However, aging was associated with an increase in correlation values for the <300 Hz range (envelope) (p = 0.034, r_e_ = 0.444, df = 21; Figure 2B, right).

Spectral amplitudes derived via a fast Fourier transform were averaged in the 105–115 Hz frequency band to assess neural synchronization to the temporal envelope (i.e., fundamental frequency; F0) of the speech-like sound. For the caudal channel, neural synchronization at the envelope frequency was larger for young compared to aged rats (p = 1.96^e-4^, r_e_ = 0.701, df = 21; Figure 2C, left). Although there was a similar trend in the rostral channel, this was not statistically significant (p = 0.069, r_e_ = 0.385, df = 21; Figure 2C, right).

Taken together, our neural synchronization measures show that for caudal generators (auditory nerve, cochlear nucleus) synchronization was either similar or increased for younger compared to aged rats, whereas for rostral generators (lateral lemniscus, inferior colliculus) synchronization was either similar or increased for aged compared to younger rats. These data may suggest an age-related relative increase in synchronization strength along the ascending auditory pathway. Extracellular recordings from inferior colliculus neurons were obtained to further explore the age-related gain increases in the auditory system to the complex, speech-like sound.

### LFP onset response and neural synchronization to the envelope are reduced in aged animals

Although changes in the neural representation of the speech-like sound were observed in the scalp-recorded synchronization measures, these EFRs reflect the superposition of activity from multiple generators. In order to localize age-related changes to the inferior colliculus and its afferents, we used LFPs as a proxy for the synaptic input to the inferior colliculus neurons. LFP time courses are displayed in Figure 3A. Neural responses to the sound onset (RMS amplitude 0–0.08 s) was larger for young compared to aged animals (p = 4.47^e-9^, r_e_ = 0.368, df = 237; Figure 3A right), suggesting that the strength of the synaptic inputs to the inferior colliculus neurons decrease with age.

**Figure 3:**
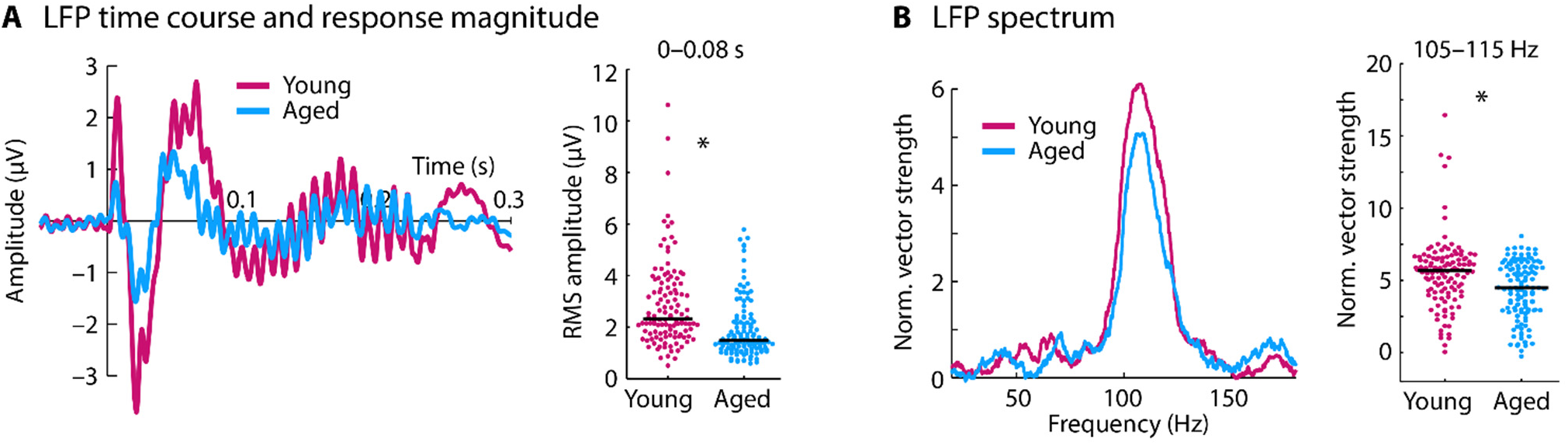
Time course and spectrum for local field potentials. **A:** Time course for each age group (left) and the root mean square (RMS) amplitude for the 0–0.08 s time window (right). **B:** Spectrum of normalized vector strength (left) and the mean vector strength for the 105–115 Hz frequency window (right). In panel A and B, each dot reflects the normalized vector strength of an individual unit. The black horizontal line reflects the median across units. *p < 0.05

In order to investigate whether this decrease in synaptic input is accompanied by a decrease in LFP synchronization, the normalized vector strength at the F0 frequency (105–115 Hz) was measured in the sustained response of the LFP (0.08–0.225 s; Figure 3B). Neural synchronization was larger for young compared to aged rats (p = 2.66^e-4^, r_e_ = 0.234, df = 237). These results indicate that the synaptic inputs to the inferior colliculus neurons in response to a speech-like sound decrease in amplitude and synchrony with age.

### Synchronization of multi-unit activity with the speech envelope are increased with age

In order to investigate the consequences of the age-related decrease in synaptic inputs (indicated by the LFPs) on the neural output of inferior colliculus neurons, multi-unit spiking activity was used as a proxy for neural population or network responses in the inferior colliculus. Time courses for the multi-unit activity (MUA) are shown in Figure 4. Spontaneous activity, quantified as the mean response in the time window preceding sound onset (–0.05–0 s), was larger in aged compared to young rats (p = 2.91^e-10^, r_e_ = 0.393, df = 237; box plots in Figure 4A, left). Unlike the onset amplitudes of the LFPs (for which responses were reduced for aged animals), there was no difference in MUA neuronal responses to the sound onset (amplitude in the 0–0.08 s time window) between age groups (p = 0.894, r_e_ = 0.009, df = 237; Figure 4A, middle and right).

**Figure 4:**
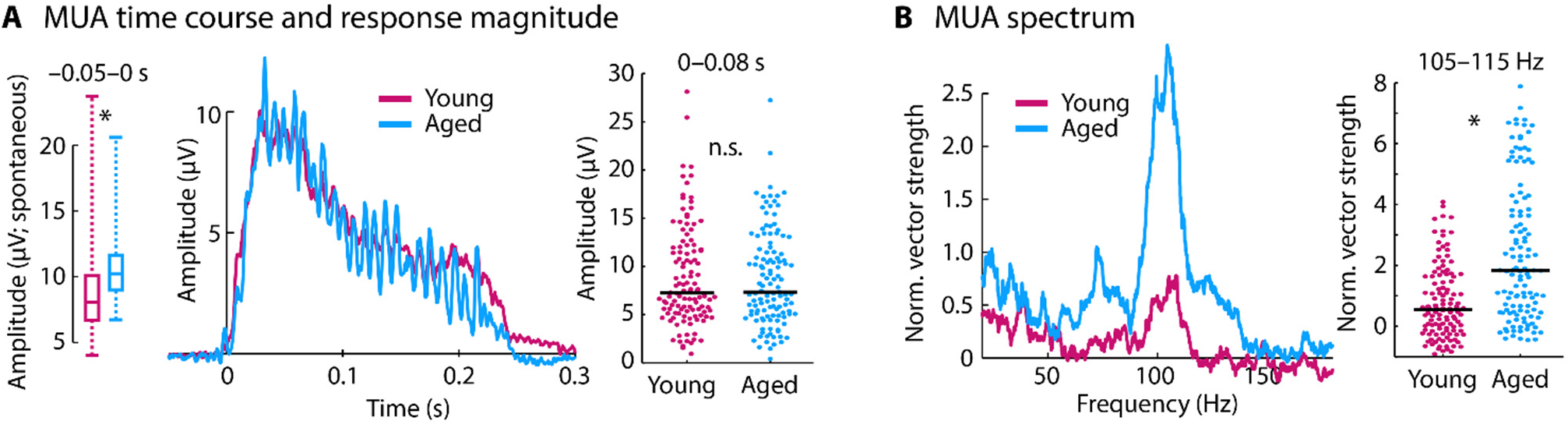
Time course and spectrum for multi-unit activity. **A:** Left: The box plots show the amplitude of the pre-stimulus onset time window (–0.05–0 s; i.e., spontaneous activity). Pre-stimulus onset activity was larger in aged compared to young animals (p < 0.05). Middle: Time course for each age group (data are baseline corrected, i.e., the mean response in the –0.05–0 s time window was subtracted from the amplitude at each time point). Right: Mean amplitude for the 0–0.08 s time window. **B:** Spectrum of normalized vector strength (left) and the mean vector strength for the 105– 115 Hz frequency window (right). In panel A and B, each dot reflects the normalized vector strength of an individual unit. The black horizontal line is the median across units. *p < 0.05

Synchronization of MUA with the envelope of the sound was quantified using normalized vector strength (Figure 4B). At the stimulation frequency (mean across the 105–115 Hz frequency window), neural synchronization was larger for aged compared to young rats (p = 4.79^e-9^, r_e_ = 0.367, df = 237). Taken together, the LFP and MUA results suggest a relative increase in neural response and an increase in synchronization from a neuron’s input to the neural population spiking output for aged compared to young animals.

### Synchronization of single-unit activity with the speech envelope are similar between young and aged animals

Neuronal activity from well isolated neurons (single-unit activity) was analyzed in order to investigate whether individual neurons in the aged inferior colliculus also show a relative enhancement of onset-evoked activity and synchronization to the envelope.

Peri-stimulus time histograms for single-unit activity (SUA) are shown in Figure 5. Spontaneous firing rates (i.e., in the –0.05–0 s time window) did not differ between age groups (p = 0.831, r_e_ = 0. 015, df = 210). Firing rates to the sound onset (0–0.08 s) were larger for young compared to aged animals (p = 1.02^e-4^, r_e_ = 0.268, df = 210; Figure 5A). Neural synchronization to the envelope (F0) of the sound measured using normalized vector strength showed no effect of age (mean across the 105–115 Hz frequency window; p = 0.703, r_e_ = 0.026, df = 210; Figure 5B). Given the reduced LFP synchronization, these results suggest an age-related increase in neural synchronization in single unit activity (a neuron’s output) relative to the LFP (synaptic input), albeit to a lesser degree than for multi-unit activity.

**Figure 5:**
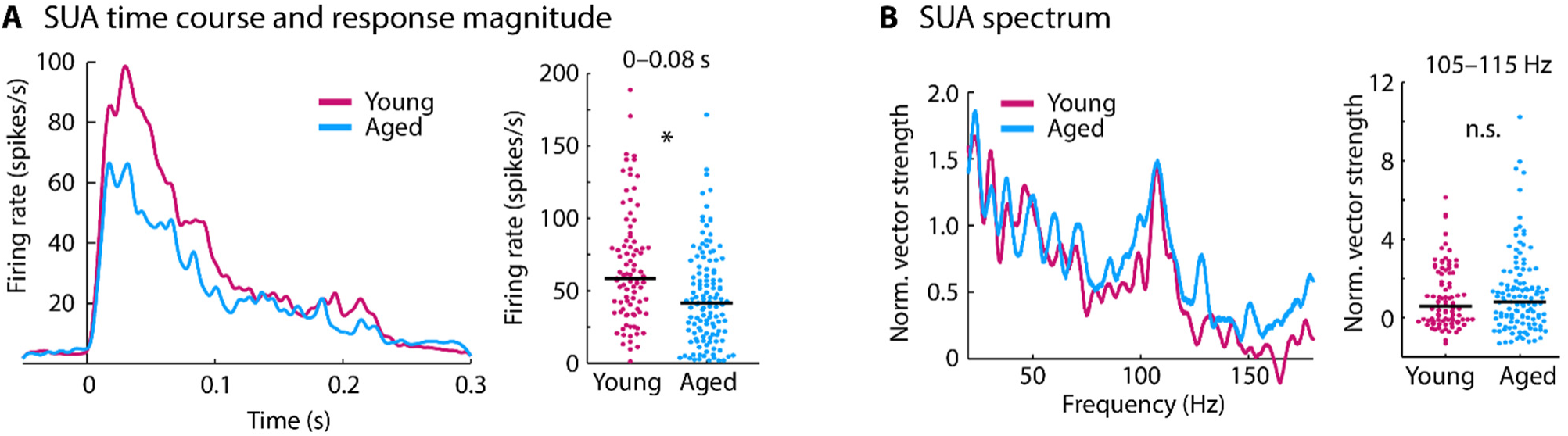
Time course and spectrum for single-unit activity. **A:** Peri-stimulus time histogram for each age group (left) and mean firing rate for the 0–0.08 s time window (right). **B:** Spectrum of normalized vector strength (left) and the mean vector strength for the 105–115 Hz frequency window (right). In panel A and B, each dot reflects the normalized vector strength of an individual unit. The black horizontal line is the median across units. *p < 0.05

### Aging affects the relative change from LFP to spiking activity

In order to quantify the effects of aging on the relative changes from the local field potential to spiking activity, we calculated the normalized difference between the LFP and SUA, and between the LFP and MUA. In detail, the LFP, SUA, and MUA were separately z-transformed by subtracting the mean and dividing by the standard deviation (including data points from young and aged rats). The z-transformation was separately calculated for onset responses (i.e., RMS or mean for the 0–0.08 s time window, for LFP and spiking activity, respectively) and neural synchronization (i.e., the mean normalized vector strength for the 105–115 Hz frequency window). For the LFP-to-SUA change, we subtracted the z-transformed LFP measure from the z-transformed SUA measure. The LFP-to-MUA change was calculating similarly by using the MUA instead of the SUA.

Wilcoxon’s rank sum test was used to test for differences between age groups (Figure 6). For onset responses, there was no difference between age groups for the relative change from LFP to SUA (p = 0.103, r_e_ = 0.115, df = 202), but a significantly larger change from LFP to MUA in aged compared to young rats (p = 1.47^e-8^, r_e_ = 0.356, df = 237). For neural synchronization, the change from LFP to SUA (p = 0.018, r_e_ = 0.167, df = 199) as well as the change from LFP to MUA (p = 7.10^e-12^, r_e_ = 0.425, df = 237) was increased for aged compared to young rats.

**Figure 6:**
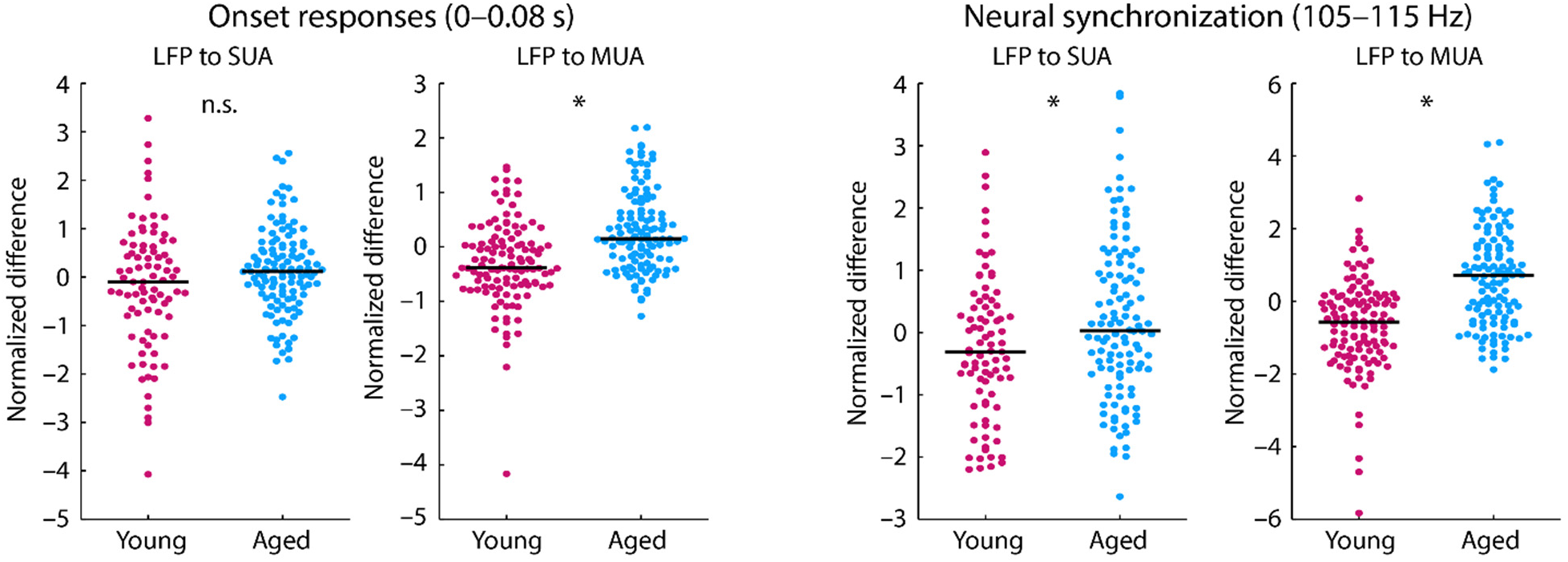
Normalized difference between local field potential and spiking activity (SUA and MUA). **Left:** Normalized difference for onset responses (time window from 0 to 0.08 s). **Right:** Normalized difference for neural synchronization in the 0.08-0.225s time window (frequency window from 105 to 115 Hz). Each dot reflects an individual unit. The black horizontal line is the median across units *p < 0.05; n.s. – not significant.

### Neural responses to a natural stimulus also show enhanced envelope sensitivity

In order to examine whether the observed effects using the noise carrier also translate to real speech, we additionally recorded neural activity in response to the original /ba/ speech sound for a subset of units. Neurons responsive to this stimulus were found in lesser numbers due to the differences in the frequency sensitivity of the rat’s hearing range. Local field potentials and multi-unit activity was recorded for 52 units in young animals and 20 units in aged animals. Single-unit activity was available for 33 units in young animals and 20 units in aged animals.

The spectrum of normalized vector strength was calculated for LFPs, MUA, and SUA. The results are displayed in Figure 7 and approximately mirror the results reported for the noise carrier. Effects of aging on neural synchronization to the stimulus envelope were assessed by comparing the mean normalized vector strength in the 105–115 Hz frequency window between age groups. For LFPs, there was no difference in neural synchronization between age groups (p = 0.730, r_e_ = 0.042, df = 70). An age-related increase in synchronization with the sound’s envelope was observed for the MUA (p = 9.78^e-3^, r_e_ = 0.303, df = 70). For SUA, there was no effect of age for neural synchronization to the envelope of the stimulus (p = 0.345, r_e_ = 0.132, df =51).

**Figure 7:**
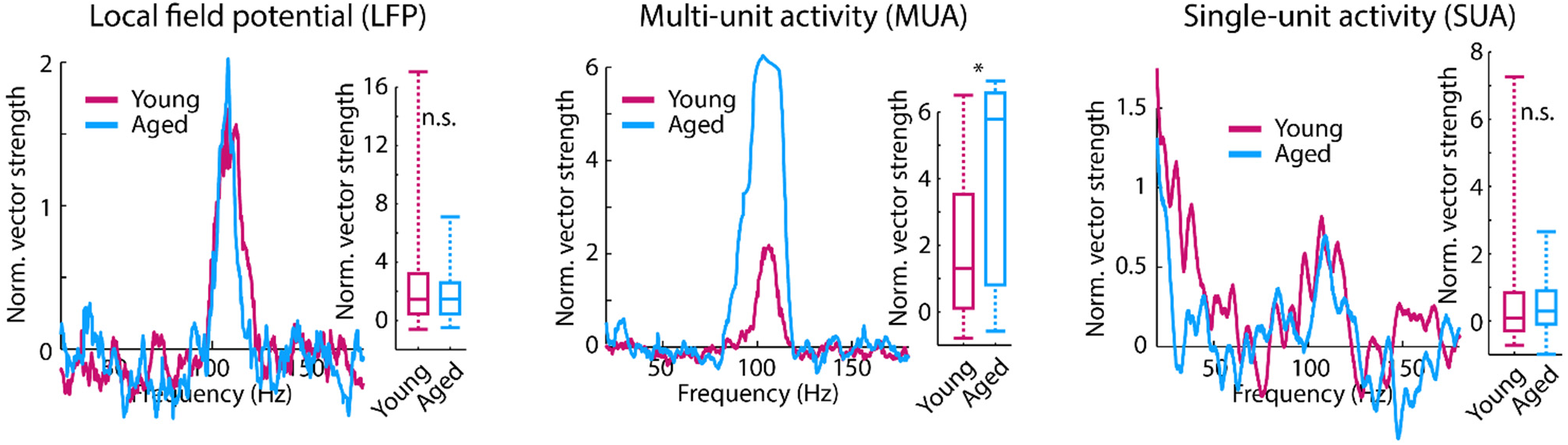
Spectrum of normalized vector strength for LFPs, MUA, and SUA in response to a speech sound. Box plots show the mean normalized vector strength for the 105–115 Hz frequency window. *p < 0.05, n.s. – not significant

## DISCUSSION

In the current study, we used a systems level (scalp recordings) and a micro-circuit level (LFPs and unit activity) approach to investigate the age-related changes in neural sensitivity to temporal regularity in a speech-like sound. Scalp-recorded potentials indicate that aging leads to a relative increase in neural synchronization to the periodicity envelope along the ascending auditory pathway. The underlying cellular changes in the midbrain were examined by recording neural activity from neurons in the inferior colliculus in response to the speech-like sound. We used the local field potential as a proxy for a neuron’s or neural population’s synaptic inputs, and multi-unit and single-unit activity as a measure of spiking output. LFP amplitudes to the sound onset and LFP neural synchronization to the temporal regularity of the envelope were smaller in aged compared to young rats. In contrast, multi-unit activity and, to a lesser degree, single unit activity showed an aged-related relative increase in synchronization to the periodicity envelope. Our results suggest that aging is associated with altered sensitivity to sounds and a sound’s temporal regularities, and that these effects may be due to altered gain in neural network activity in the aged auditory midbrain.

### Scalp-evoked potentials point to a decrease in brainstem responses and an increase in inferior colliculus responses

Auditory brainstem responses (ABRs) and neural synchronization (i.e., EFRs) have been used to study age-related changes in neural activity to speech and other complex sounds in humans (Stamper and Johnson, 2015, Clinard and Tremblay, 2013, Anderson et al., 2012). The ability to obtain these responses non-invasively makes them an ideal bridge between human studies and studies in animal models, for which the underlying pathophysiology can additionally be studied at the micro-circuit level using more invasive techniques (Zhong et al., 2014, Shaheen et al., 2015).

Consistent with previous studies (Fernandez et al., 2015, Sergeyenko et al., 2013, Parthasarathy et al., 2014), we observed an age-related decrease in the wave 1 amplitude of the ABR to clicks at suprathreshold sound levels (Figure 2A). The wave 1 of the ABR at suprathreshold sound levels may be considered as a physiological indicator for the degree of cochlear synaptopathy, that is, the loss of synapses between the inner hair cells and the auditory nerve fibers (Kujawa and Liberman, 2009, Konrad-Martin et al., 2012). This cochlear synaptopathy has been observed in aging and noise-exposed animals, including mice (Sergeyenko et. al., 2013, Fernandez et. al., 2015), rats (Mohrle et. al., 2016), gerbils (Gleich et. al., 2016), guinea pigs (Furman et. al., 2013, Schmeidt et. al., 1996), and macaques (Valero et. al., 2017), in addition to evidence from post-mortem human temporal bones (Viana et. al., 2015). The reduction of wave 1 in aged compared to young rats thus suggests the presence of cochlear synaptopathy in the aged population used in the current study.

The sensitivity of the scalp-recorded EFRs to regular temporal structure in a speech-like sound was investigated using measures of neural synchronization. We observed reduced synchronization to the envelope and fine structure information for activity likely originating in the auditory nerve and cochlear nucleus (Figure 2, left). Signals likely originating from rostral generators, including the inferior colliculus, showed an age-related increase in neural synchronization to the envelope of the speech-like sound (Figure 2B, C). These results suggest an age-related transformation in the neural representation of envelope cues in speech along the ascending auditory pathway, which we investigated further using more invasive extracellular electrophysiological methods.

### Aging increases network level activity of inferior colliculus neurons relative to their synaptic inputs

In order to study the contributions of the auditory midbrain to the changes seen in the scalp-recorded responses, we simultaneously recorded LFPs and unit responses from the same site in response to sound. LFPs are thought to be largely the aggregate pre-synaptic activity at the dendrites and the soma, and hence a proxy for the inputs to the neuron or neuronal region (Buzsaki et al., 2012, Gourevitch and Edeline, 2011, Logothetis and Wandell, 2004, Logothetis et al., 2001). By comparing these LFPs to the spiking output, the transformation of these sound representations in the inferior colliculus can be investigated (Herrmann et al., 2017).

In the current study, LFPs to the sound onset and neural synchronization to the sound’s envelope were decreased for aged compared to young rats (Figure 3), suggesting that inputs to the inferior colliculus neurons are degraded. In contrast, multi-unit spiking activity in the inferior colliculus – reflecting population or network output – revealed no age difference in the onset response and age-related increase in synchronization to the sound’s envelope (Figure 4). This relative enhancement seen at the population level (MUA) was also present, albeit to a lesser degree, at the level of the single neurons (SUA, Figures 5 and 6). These results suggest that the inferior colliculus neurons selectively increase their neuronal activity to the envelope of a speech-like sound in aged animals, despite the reduced synaptic inputs to these neurons.

Previous work suggested that midbrain neurons responding to simple stimuli like tones and noise bursts show remarkably subtle changes with age (Willott et al., 1988a, Willott et al., 1988b). This is despite the extensive degradation of the peripheral auditory system, such as the loss of outer hair cells (Chen et al., 2009, Spongr et al., 1997), changes in endocochlear potential (Ohlemiller et al., 2006), and the loss of cochlear synapses (Sergeyenko et al., 2013) and spiral ganglion neurons in the auditory nerve (Bao and Ohlemiller, 2010). Even when studies using single-unit recordings do find age-related changes in temporal processing (Palombi et al., 2001, Walton et al., 2002), it is unclear whether these changes are inherited from previous stages of auditory processing, or generated in the nucleus being studied. Our approach in this study shows that the enhancements seen in the spiking output of the inferior colliculus neurons, in particular at the population and network level, may explain some of this dichotomy in peripheral versus central responses with age.

### Potential mechanisms underlying age-related changes in sensitivity to temporal regularity in sounds

Acute insults (e.g., ototoxic drugs or mechanical deafening) to the auditory system damages the auditory periphery and triggers homeostatic changes in the auditory pathway (Kotak et al., 2005, Chambers et al., 2016). The most drastic change reported is a loss of inhibitory circuits and function (Resnik and Polley, 2017, Caspary et al., 2013, Richardson et al., 2013, Ling et al., 2005), which, in turn, increases neuronal activity in the central auditory structures (Chambers et al., 2016). The current data in combination with recent evidence suggests that central auditory systems undergo similar gain increases when the peripheral insult is gradual as is the case for aging (Lai et al., 2017, Parthasarathy et al., 2014, Presacco et al., 2016).

The increase in neuronal activity due to a loss of inhibition is typically accompanied by a reduction in the precision of neural coding. Inferior colliculus neurons change their tuning curves to become less selective with age (Leong et al., 2011, Rabang et al., 2012). In addition, sensitivity to temporal regularities in simple stimuli increases for slow amplitude modulations but decreases for fast amplitude modulations in aged animals (Walton et al., 2002, Herrmann et al., 2017). In the current study, inferior colliculus neurons in aged animals showed an increase in the spontaneous firing rate (Figure 4A), which suggests inhibition was reduced in the aged midbrain. Furthermore, despite reduced synaptic inputs to inferior colliculus neurons (as indexed by our LFP recordings), neuronal firing of populations of inferior colliculus neurons was overly synchronized with the envelope of a speech-like sound in aged compared to young animals (Figures 3 and 4). Our data thus suggest that temporal response precision is altered in the aged midbrain due to hypersensitivity in networks of neurons.

Increased gain in the auditory system may lead to better detection of weak signals in neural circuits, and may, in turn, support hearing in quiet environments (Chambers et. al., 2016). For example, hearing loss has been shown to increase sensitivity to suprathreshold amplitude modulation in sounds (Moore et al. 1996). However, neural gain enhancements may come at the cost of poor discrimination between stimuli (Guo et al., 2017). For example, discrimination between fundamental frequency in speech is reduced for older adults with clinically normal audiograms (Vongpoisal and Pichora-Fuller, 2007), and a gain increase in cortex has been linked to poor speech perception in the presence of competing sound (Millman et al., 2017). Hence, an aberrant neural gain increase in aging may not be beneficial for listening to sounds that are typical of everyday life, such as speech, when perception requires the segregation of different concurrent sounds.

## CONCLUSIONS

We investigated how aging affects neural sensitivity to temporal regularity in a speech-like sound. Systems level recordings (i.e., at the scalp) were combined with microcircuit recordings (i.e., LFPs and unit activity recorded extracellularly) in young and aged rats. We show that aging is associated with increased neural activity along the ascending auditory pathway, which alters the sensitivity of midbrain (i.e., inferior colliculus) neurons to temporally regular structure in sounds. Specifically, synchronization to the periodicity envelope in speech (at the fundamental frequency) was enhanced for spiking activity of populations of inferior colliculus neurons in aged rats, despite the neuron’s reduced synaptic input. The data suggest that temporal response precision is altered in the aged midbrain due to hypersensitivity in networks of neurons.

## ACKNOWLEDGEMENTS

This study was supported by NIDCD DC-011580 to ELB.

